# Characterization of nitrogenase genes *nifP* and *nifVZT1* in the cyanobacterium *Anabaena* (*Trichormus*) *variabilis* ATCC 29413

**DOI:** 10.1101/2025.04.01.646610

**Authors:** Brenda Pratte, Susan Vernon, Teresa Thiel

## Abstract

*Anabaena (Trichormus) variabilis* ATCC 29413 is a heterocyst-forming cyanobacterium with two Mo-nitrogenases encoded by the large *nif1* and *nif2* gene clusters. The *nif1*-encoded nitrogenase is expressed in heterocysts under oxic growth conditions, whereas the nif2-encoded nitrogenase is expressed only in anoxic vegetative cells. The *nifP* and *nifVZT1* genes are contiguous and distinct from the major *nif1* and *nif2* clusters, suggesting that *nifP* and *nifVZT1* may be cotranscribed. However, we identified primary transcription start sites for *nifP* and *nifVZT1*, indicating that each has its own promoter. The promoter region of *nifP* shares multiple conserved motifs with the *nifB1* and *nifB2* promoters, which are regulated by homologous transcriptional activators CnfR1 and CnfR2, respectively. These conserved motifs are not present in the promoter region of *nifVZT1*. Using *nifP* promoter fragments that included or excluded the conserved motifs fused to reporter gene *lacZ*, we determined which motifs contributed to *nifP* expression; however, these identified motifs differed from the conserved motifs shown previously to regulate *nifB1* or *nifB2* transcription. Further, the regulation of transcription requires a region inside the coding region of *nifP*. Mutant strains lacking *nifP* or *nifVZT1* grew well in the absence of fixed nitrogen and showed no reduction in nitrogenase activity compared to the wild-type strain grown under oxic or anoxic conditions. Further, *nifP* was expressed constitutively in vegetative cells grown −N or +N, but its expression increased greatly in heterocysts by CnfR1 activation of the *nifP* promoter. In contrast, *nifVZT1* was expressed only in heterocysts but, unlike most other *nif1* genes, was not under the control of CnfR1.

## Introduction

Nitrogen fixation in cyanobacteria occurs in both unicellular and filamentous strains; however, under aerobic growth conditions, it is largely restricted to strains that can differentiate cells called heterocysts that develop in response to nitrogen deficiency (Herrero et al., 2013; Kumar et al., 2010; Wolk et al., 1999; Zhang et al., 2006).

Heterocysts form in a semi-regular pattern in a filament comprising 5–10% of the cells. Their structure and metabolism are designed to protect the enzyme nitrogenase from inactivation by oxygen. Thus, heterocysts inactivate photosystem II, which produces oxygen; have a glycolipid layer that restricts oxygen diffusion in the cell; and have high levels of respiration to consume intracellular oxygen (Murry et al., 1984; Murry and Wolk, 1989; Valladares et al., 2003; Walsby, 1985, 2007).

*Anabaena (Trichormus) variabilis* ATCC 29413 is a heterocyst-forming cyanobacterium with three nitrogenases that are expressed in different environments (reviewed in (Thiel, 2004; Thiel and Pratte, 2013; Thiel and Pratte, 2014)). There are two Mo-nitrogenases encoded by the *nif1* and *nif2* genes and a V-nitrogenase encoded by the *vnf* genes. The Nif1 Mo-nitrogenase and the V-nitrogenase are heterocyst specific, while the Nif2 Mo-nitrogenase is expressed only under anoxic conditions in vegetative cells. All three nitrogenases are made only in the absence of fixed nitrogen, and the V-nitrogenase is made only when Mo concentrations are very low. Although Mo does not regulate the expression of the *nif1* or *nif2* genes, it is required for the synthesis of a functional Mo-nitrogenase. Similarly, V does not regulate the *vnf* genes but is required for the synthesis of a functional V-nitrogenase. The Nif1 Mo-nitrogenase and the V-nitrogenase are made late in heterocyst development under appropriate conditions of Mo availability about 24 h after deprivation of fixed nitrogen, ensuring that nitrogenase is protected from oxygen. The Nif2 nitrogenase is made only in anoxic vegetative cells by about 6 h after nitrogen removal.

In cyanobacteria, the primary regulator of the *nif* genes is CnfR, an activator protein with two N-terminal 4Fe-4S binding sites, similar to bacterial ferredoxins, and a C-terminal DNA-binding domain (Jones et al., 2003). In *Nostoc* sp. PCC 7120, a *cnfR* (formerly called *patB*) mutant has very low levels of nitrogenase and does not grow diazotrophically (Jones et al., 2003). Similarly, the *nif* genes of the filamentous non-heterocystous cyanobacterium *Leptolyngbya boryana* are not expressed in a *cnfR* deletion mutant (Tsujimoto et al., 2014; Tsujimoto et al., 2016). *A. variabilis*, with two *nif* clusters, has a *cnfR* gene to regulate each cluster. Deletion of *cnfR1* abolishes the activation of *nifB1* in heterocysts, while loss of *cnfR2* prevents the expression of *nifB2* in anaerobic vegetative cells (Pratte and Thiel, 2016, 2021; Thiel, 2019). The *vnf* genes in *A. variabilis* are regulated by dual repressors, VnfR1 and VnfR2 (Pratte et al., 2013).

In *A. variabilis*, the contiguous *nif1* genes (*nifB_1_S_1_U_1_H_1_D_1_K_1_E_1_N_1_X_1_W_1_*)and the contiguous *nif2* genes (*nifB_2_S_2_U_2_H_2_D_2_K_2_E_2_N_2_X_2_W_2_)* are transcribed primarily from the promoters for the first gene in each cluster, *nifB1 or nifB2* (Pratte and Thiel, 2014; Thiel, 2019; Ungerer et al., 2010; Vernon et al., 2017). In *A. variabilis*, the single copy of *nifP* is not contiguous with either the *nif1* or *nif2* gene cluster. *L. boryana* has two divergently transcribed operons, *nifBSUHDKVZT* and *nifPENXW,* with a shared promoter region between *nifB* and *nifP* (Tsujimoto, et al., 2016). The *nifB1*, *nifB2*, and *nifP* promoter regions in *A. variabilis* (Vernon et al., 2017) and the *nifB-nifP* intergenic region in *L. boryana* (Tsujimoto et al., 2016) have multiple well-conserved motifs whose losses yield variable effects on promoter activity. Loss of either of the first two of three repeated 8-bp TGAGTACAA motifs in *A. variabilis* results in a similar 50% decrease in transcription of *nifB1* with a further decrease when both motifs are mutated (Vernon et al., 2017).

Although the conserved regions of *nifB1* and *nifB2* in *A. variabilis* are similar, the replacement of the *nifB2* promoter region with the similar region from *nifB1* results in a complete loss of *nifB2* expression (Vernon et al., 2017). In addition to the conserved *nifB2* 5′ untranslated region, expression of *nifB2* requires sequences within the 5′ end of the *nifB2* gene (Vernon et al., 2017).

Nitrogenase is a highly conserved complex of two distinct enzymes, dinitrogenase and dinitrogenase reductase, whose structural properties have been elucidated for the enzyme from the proteobacterium *Azotobacter vinelandii*. Dinitrogenase reductase (encoded by *nifH*) accepts electrons from donors such as NADH and transfers them to dinitrogenase (Lawson and Smith, 2002). Dinitrogenase comprises two α-subunits (encoded by *nifD*) and two β-subunits (encoded by *nifK*), which form a heterotetramer that includes two active site FeMo-cofactors [MoFe_7_S_9_C-homocitrate] (Einsle, 2014) and two P-clusters [8Fe:7S] (Peters et al., 1997). These dinitrogenase metallo clusters accept electrons from the [Fe_4_-S_4_] cluster that is part of the dinitrogenase reductase dimer (reviewed in (Einsle and Rees, 2020; Rubio and Ludden, 2008; Schwarz et al., 2009). In addition to the major structural proteins NifH, NifD, and NifK, NifB, and NifEN are required for FeMo-co synthesis and assembly of the active nitrogenase complex.

Further, NifH plays important roles in both P-cluster formation and metalloprotein complex formation (Buren et al., 2020). Although these proteins are sufficient for the enzyme complex, a number of ancillary proteins are required to produce the metalloclusters, which are essential for nitrogenase activity and serve as carrier proteins (Ribbe et al., 2014; Rubio and Ludden, 2008). In addition to these metalloclusters, pathways to provide homocitrate, S-adenosyl-methionine, and Mo (or V) for the complexes are also required (Buren et al., 2020). Further details of the synthesis and the functions of all these *nif*-related components are provided in a comprehensive review of the proteins and factors required for the assembly of the active Mo-nitrogenase in *A. vinelandii* (Buren et al., 2020). However, a number of the proteins present in *A. vinelandii*, including homologs of NifQ, the Mo donor to FeMo-co (Hernandez et al., 2008), NifM, which stabilizes NifH (Howard et al., 1986), and NafY, which delivers FeMo-co to the NifDK nitrogenase complex (Hernandez et al., 2011; Phillips et al., 2021) are absent in cyanobacterial genomes.

Here, we characterize the ancillary cyanobacterial nitrogenase genes *nifP*, *nifV1*, *nifZ1* and *nifT1*. NifP (CysE) is a serine O-acetyltransferase (Evans et al., 1991). NifV synthesizes homocitrate, a component of FeMo-cofactor (Rubio and Ludden, 2008), from acetyl-coenzyme A and 2-oxoglutarate. NifZ, with NifH, aids in P-cluster assembly and maturation (Hu et al., 2004), and NifT is a small, conserved, nonessential protein of unknown function (Beynon et al., 1988; Stricker et al., 1997). Associated with the two *nif* clusters in *A. variabilis* are two copies of *nifZ* and *nifT*. One copy of *nifZ* and *nifT* is associated with *nifV1,* while the second copy is associated with the large *nif2* cluster (see **Fig. 1A**). In contrast, there is a single copy of *nifP* and three copies of *nifV*. We characterized the expression of *nifP* and *nifVZT1* genes and determined whether they are required for nitrogenase activity in *A. variabilis*.

**Fig. 1.**
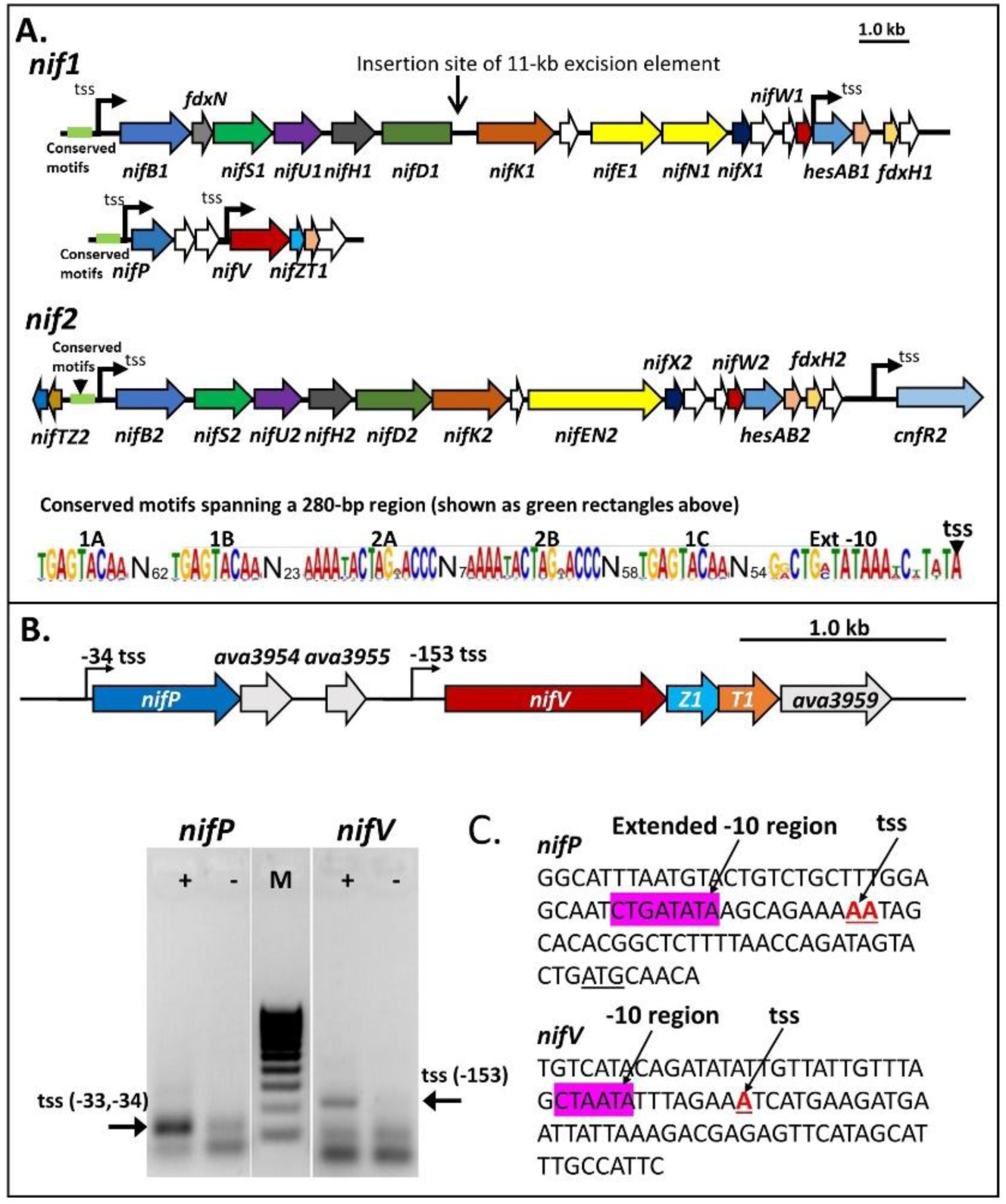
Gene organization of *nifP* and *nifVZT1* and transcription start sites. **A**. Maps of *nif1* and *nif2* gene clusters, and *nifP nifVZT1* with the conserved motifs upstream of *nifB1*, *nifB2*, *nifZ2*, and *nifP*. B. The genome region with *nifP* and *nifVZT1* genes and the sequenced PCR products for the transcription start sites using 5’ RACE with (+) or without (-) RNA 5′ polyphosphatase. M = MW markers. **C**. Location of transcription start sites (tss) for *nifP* and *nifV* and possible -10 regions. The translation start site for *nifP* is underlined in black.

## Results

### Unique primary transcription start sites for *nifP* and *nifVZT1*

The *nifP* gene is not part of the large *nif* operon in cyanobacteria; it is upstream of the *nifVZT1* operon; however, the *nifP* promoter region has the conserved elements found upstream of *nifB1* and between the divergently transcribed *nifB2* and *nifZ2T2* genes (Fig. 1A). We showed previously that the *nifB1* promoter drives transcription of the large *nif1* gene cluster in *A. variabilis* and that the apparent strong promoter for *nifH1* represents a transcriptional processing site (Ungerer et al., 2010). To determine whether *nifP* and *nifVZT1* are under the control of the same promoter, we determined the transcription start sites for both genes by 5’ RACE (rapid amplification of cDNA ends), a method that can distinguish between processed and primary transcripts (Bensing et al., 1996). Thus, we identified the 5’ end of each transcript by ligating an RNA adapter to the 5′ end of the transcript. A primary transcript requires treatment of the sample with RNA 5′ polyphosphatase to hydrolyze the 5′ triphosphate to a monophosphate for ligation. Since processed transcripts already have a 5′ monophosphate, RNA 5′ polyphosphatase is not required for ligation to the adapter. Sequencing the *nifP* RNA ligase-mediated RT-PCR product produced two products, each about 50%, corresponding to transcription start nucleotides at −33 and −34 from the translations start site (**Fig. 1B**). Sequencing of the *nifV* cDNA product yielded one clear transcription start site at nucleotide −153 from the translation start site (**Fig. 1B**). Since RNA 5′ polyphosphatase was required for good ligation of the RNA adapter, both are primary transcripts; hence, *nifP* and *nifVZT1* have their own promoters.

### Characterization of upstream activator regions for *nifP* using *nifP*:*lacZ* fusions

We and others (Brenes-Alvarez et al., 2019; Thiel, 2019; Tsujimoto et al., 2016; Vernon et al., 2017) have identified conserved motifs (**Fig. 1A**) upstream of the promoter that are important for the expression of *nifB*, and they are conserved in the region upstream of the *nifP* promoter. In order to determine the role in the expression of *nifP*, we transcriptionally fused a promoterless *lacZ* to regions upstream of *nifP* and measured β-galactosidase activity in cells grown with or without fixed nitrogen. A region that extended more than 1000 bp upstream of *nifP* in strain JJ154 had negligible activity, while a variety of smaller fragments gave only low activity (**Fig. 2A**). Based on our prior experience with *nifB2*, which requires the addition of the 5′ end of the *nifB2* coding region for good expression, we added the first 162 bp of *nifP* to the *lacZ* fusions, which not only increased *lacZ* expression of the 1031-bp (SM24) vs. JJ154 but also provided repression by fixed nitrogen (**Fig. 2B**). BP1321, which has *lacZ* integrated into the chromosome and driven by the entire upstream region of *nifP,* had *lacZ* expression and nitrogen regulation similar to SM24 (**Fig. 6**). SM4, which has a short promoter fragment with only the 1C motif, had poor activity and was not regulated by nitrogen. Incremental increases in promoter region size improved activity and provided nitrogen regulation; however, β-galactosidase activity overall increased only slightly with the addition of the 2A motif (SM5) and the 1B motif (SM6) and decreased when the region extended beyond the 1B motif (SM7) and included motif 1A (SM8). However, the increase in β-galactosidase activity with the 1B motif occurred only under –N conditions, suggesting that 1B is important for optimal expression –N. For the four strains with good β-galactosidase expression (SM18, SM5, SM6, and SM7), fixed nitrogen reduced expression by about 40-60%. The two promoter fragments driving *lacZ* in SM28 vs. SM18 differed in length by only 28 bp; however, the larger fragment greatly increased expression, especially in cells gown with fixed nitrogen (**Fig. 2B**). The sequence between motifs 1B and 1C is also conserved in the *nifP* upstream regions of other cyanobacteria and includes two partially redundant A/C-rich motifs, 2A and 2B (**Fig. 2C**). Inclusion of the 2B motif had the greatest effect in increasing *lacZ* expression; however, but 2A (SM5) and 1B (SM6) also improved expression. SM28 showed the strongest repression by fixed nitrogen. As the promoter size increased, there was greater expression +N, indicating loss of regulation in vegetative cells. Assuming that much of the increase in expression under –N conditions was attributable to vegetative cells, then the expression in heterocysts did not increase much from the shorter SM18 to the longer SM5 fragment. For example, if the activity +N for each sample is subtracted from the value –N, then the –N values for SM 28, 18, and 5 are similar. However, the inclusion of the region upstream of 1B, especially the 1A motif, decreased expression +N and –N. Thus, while the 1A, 1B, and 1C motifs have a role in the expression of *nifP*, motifs 1B and 2B are particularly important for the transcription of *nifP* under nitrogen-limiting conditions.

**Fig. 2.**
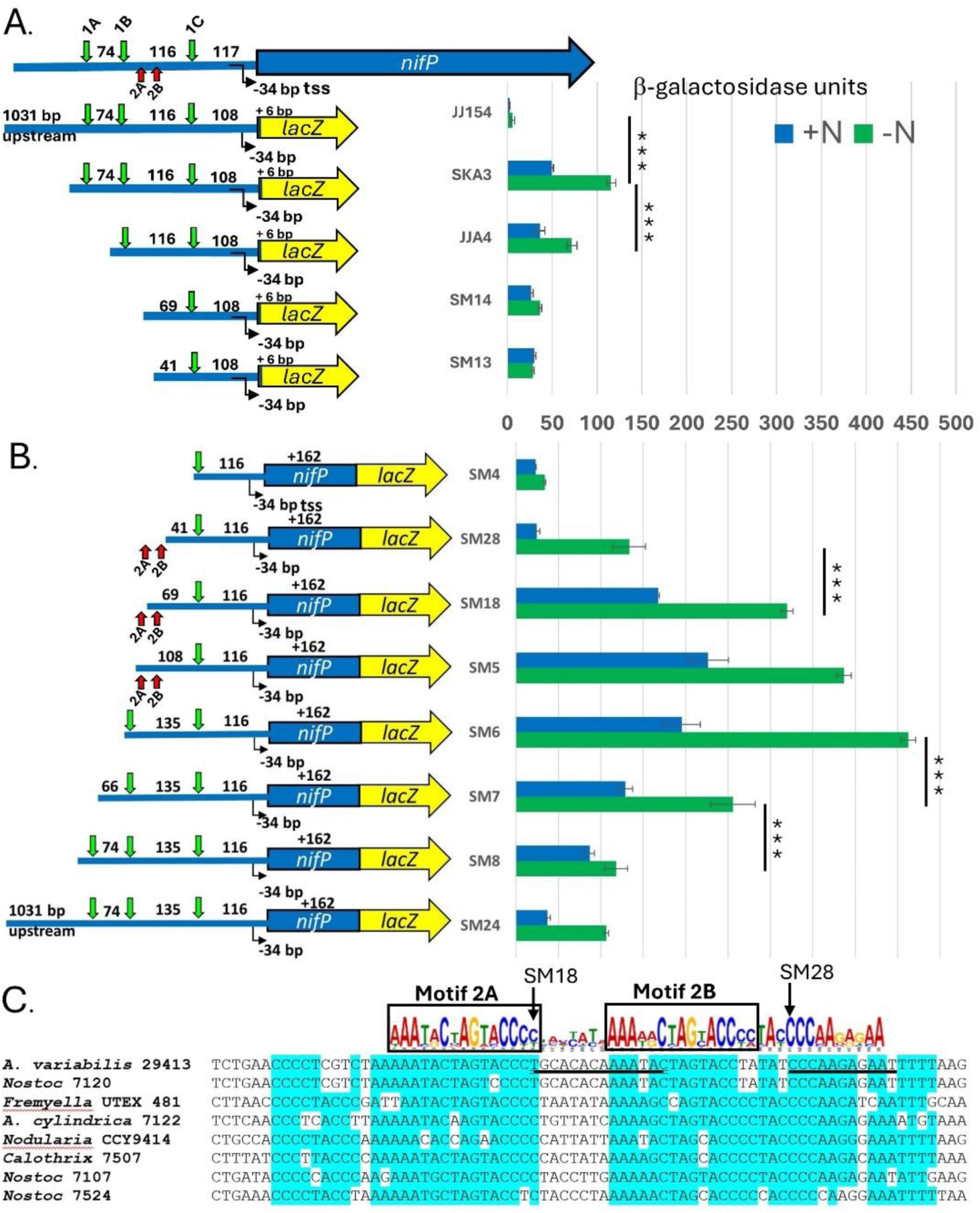
Expression of *nifP*:*lacZ* fusions. **A.** Fusions of *nifP*:*lacZ* extending only 6 bp into the *nifP* coding region. **B.** Fusion with the first 162 bp of *nifP*. Strains lacking nitrogen (-N) were grown aerobically for 24 h after nitrogen stepdown for heterocyst development. Green vertical arrows represent conserved sequences 1A, 1B, 1C (TGAGTACA) shown in Fig. 1A. Red vertical arrows represent conserved motifs 2A and 2B shown in Fig. 1A and panel C. The number of bp between the conserved sequences 1A, 1B, and 1C includes the element to the left of the number. ***, p < 0.001 (shown only for the most relevant pairs). **C.** Sequence of the region between 1B and 1C including motifs 2A and an extended version of motif 2B shown in Fig. 1A. The motifs were identified by MEME software using the *nifP* upstream sequences shown in the alignment. The primers for the 5’ ends of SM18 and SM28 are underlined in the sequence for *A. variabilis*, indicating that motif 2B is missing in SM28.

To identify the region within the *nifP* coding region that was required for transcription of the *nifP*:*lacZ* fusion, we created fusions that included the 2B motif but also extended various distances into *nifP*. Extensions of 36 bp (SM16), 66 bp (SM20), or 105 bp (SM21) into *nifP* greatly increased activity compared to the control SM14, which extends only 6 bp into *nifP* (**Fig. 3**); however, the 162-bp extension into *nifP* (SM18) showed the best repression under nitrogen replete conditions, which was also seen in other promoter constructs that extended 162-bp into *nifP* (**Fig. 2B**).

**Fig. 3.**
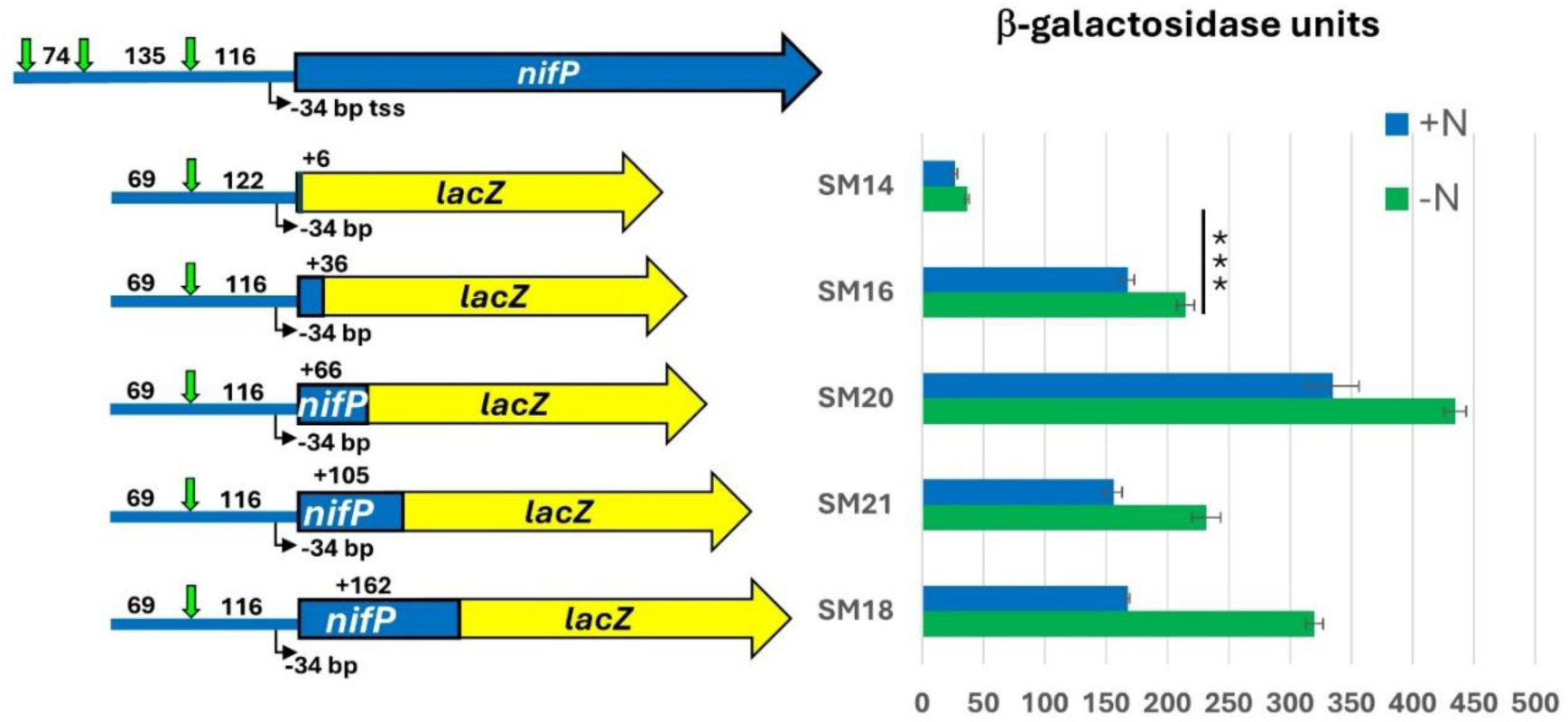
Expression of *nifP*:*lacZ* fusions. Strains lacking nitrogen (-N) were grown aerobically for 24 h after nitrogen stepdown for heterocyst development. β-galactosidase activity was determined for quadruplicate samples.***, p < 0.001 (shown only for the most relevant pair).

### Characterization of *nifP* and *nifVZT1* and *nifZT1* mutants

To determine the role of NifP in nitrogen fixation, we constructed a fully segregated deletion mutant of *nifP* (SM19) and measured its growth and nitrogenase activity compared to the wild-type (WT) strain FD. Because there are two nitrogenases in this strain, we determined whether the mutant affected either nitrogenase. Under aerobic growth conditions in the absence of fixed nitrogen when only the *nif1* genes are expressed, there was no difference in growth rate or nitrogenase activity (**Fig. 4**). Similarly, after 6 h under anoxic conditions in the absence of fixed nitrogen when only the *nif2* genes are expressed, there was no difference in nitrogenase activity (**Fig. 4A**). Growth rate could not be determined under anoxic conditions because the WT strain grows very poorly under this condition. Thus, NifP is not required for nitrogen fixation using either the Nif1 or the Nif2 nitrogenase in *A. variabilis*. Loss of *nifVZT1* (mutant BP1317) had no effect on growth or acetylene reduction (nmoles ethylene/OD_720_/min: WT = 1.74 ± 0.01; *nifVZT1* = 1.62 ± 0.73, p= 0.14). Similarly, loss of *nifZT1* (BP1318) (2.30 ± 0.55 nmoles ethylene/OD_720_/min, p= 0.09) did not impair nitrogenase activity.

**Fig. 4.**
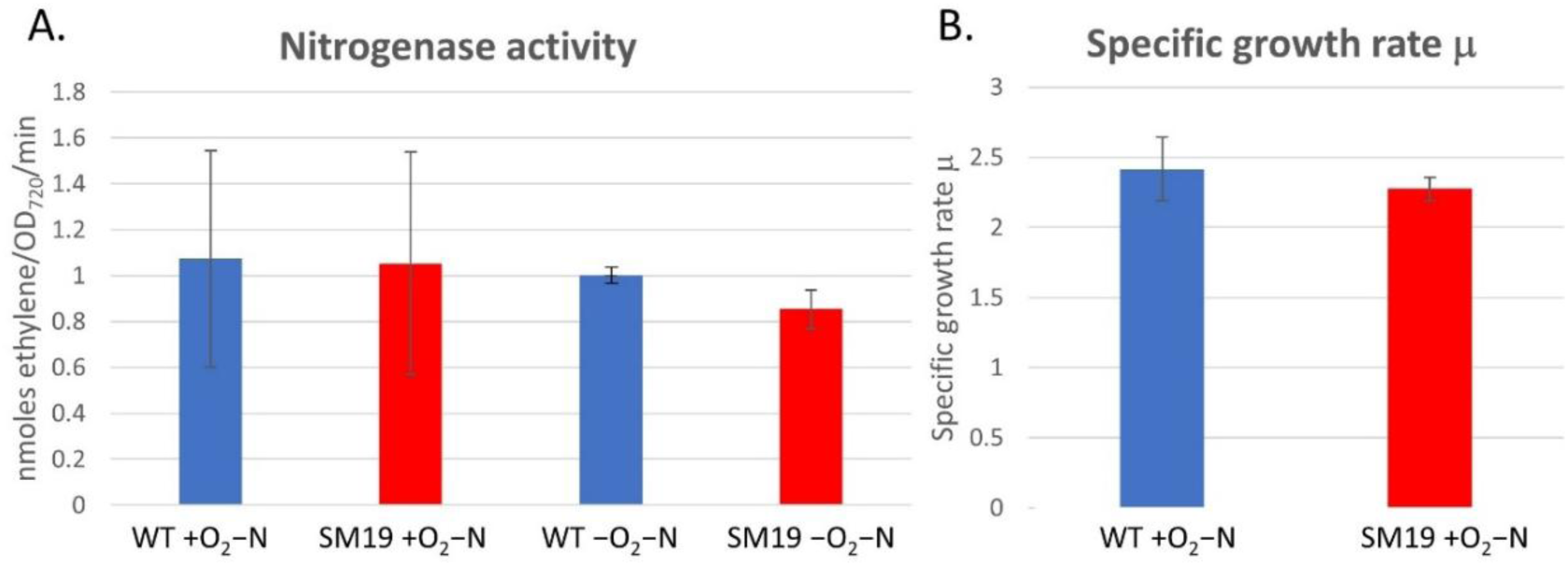
Nitrogenase activity and growth of a *nifP* mutant. Nitrogenase activity **(A)** and specific growth rate **(B)** were determined in the WT strain (FD) and the *nifP* mutant (SM19) grown without fixed nitrogen either aerobically for 24 h for heterocyst development or anaerobically for 6 h for anoxic vegetative cells. Unlabeled values were not statistically significant.

### The role of regulators CnfR1 and CnfR2 in the expression of *nifP*

Although *nifP* was not required for nitrogen fixation, the presence of the conserved sequences upstream of *nifP*, *nifB1,* and *nifB2* suggests that the genes are similarly regulated. In *A. variabilis*, CnfR1 is required for activation of the *nifB1* promoter in heterocysts, and CnfR2 is required for activation of the *nifB2* promoter in anoxic vegetative cells (Pratte and Thiel, 2016). Therefore, we measured the transcription of *nifP* in the WT strain and in *cnfR1* and *cnfR2* mutants. In addition, we measured *nifP* transcripts in cells grown −N when *cnfR1*/*cnfR2* genes are expressed and +N when *cnfR1*/*cnfR2* genes are not expressed. Expression of *nifP* in the WT strain under aerobic conditions was about 3-fold higher −N vs. +N and about 2-fold higher under anoxic conditions −N vs. +N (**Fig. 5A**), similar to our previous results (Pratte and Thiel, 2016). In the *cnfR1* mutant, under aerobic conditions −N, *nifP* expression was about half of the WT, similar to previous results with a *cnfR1* mutant that was not a deletion (Pratte and Thiel, 2016) (**Fig. 5A**). The P*_nifP_*:*lacZ* fusion had about 20% activity in a *cnfR1* mutant compared to the wild-type strain (**Fig. 6**). However, these values do not consider the fact that CnfR1 is made only in heterocysts, about 10% of the cells. Thus, if loss of CnfR1 prevented the expression of *nifP* only in heterocysts, then the two-fold loss in *nifP* expression in the *cnfR1* mutant compared to the WT strain represents a 20-fold loss in *nifP* expression in heterocysts. In a *cnfR2* mutant, under anoxic conditions −N, *nifP* expression was only slightly lower than the WT (**Fig. 5A**). Since CnfR2 is made in all anoxic vegetative cells grown −N, CnfR2 does not control the expression of *nifP*. Nothing is known about the expression of *nifV1* in anoxic cells grown −N or the role of CnfR1 or CnfR2 in the expression of *nifV1*. Therefore, we measured the transcription of *nifV1* under anoxic conditions in the WT strain and in *cnfR1* and *cnfR2* mutants.

**Fig. 5.**
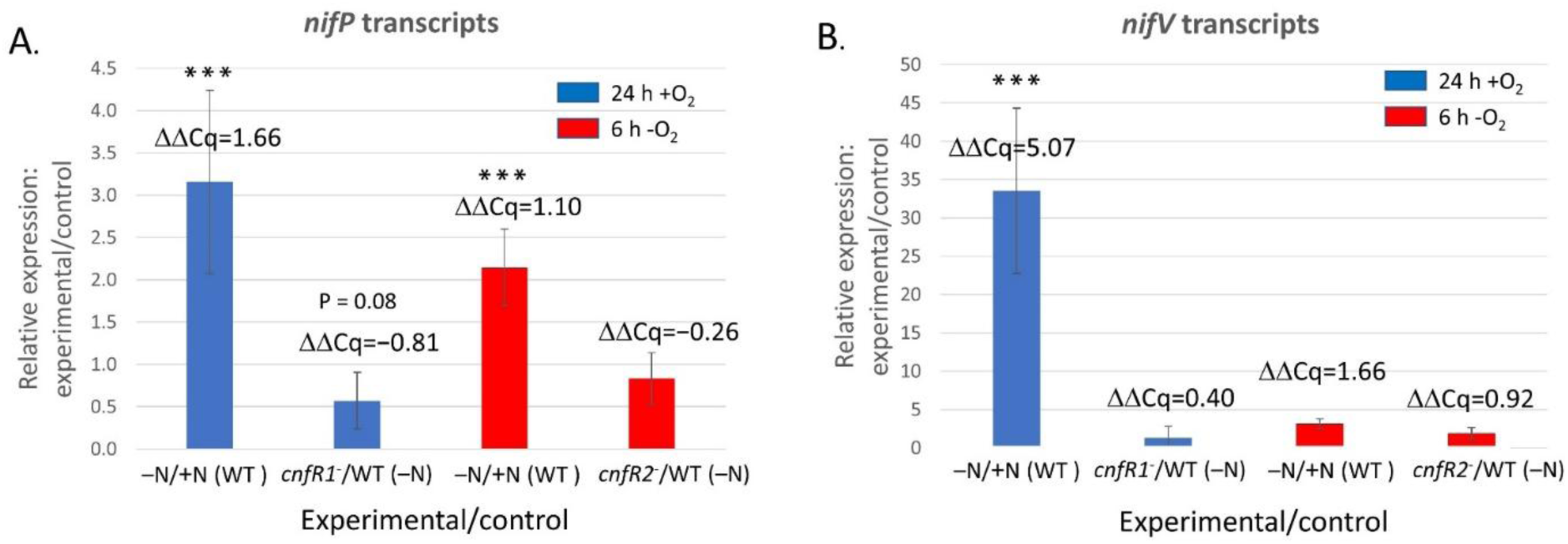
Expression of *nifP* and *nifV*. A. Expression of *nifP* (**A**) or *nifV* (**B**) was determined in the WT strain (FD), the *cnfR1* mutant strain (BP851), or the *cnfR2* mutant strain (JU802) grown with or without fixed nitrogen either aerobically for 24 h for heterocyst development or anaerobically for 6 h for anoxic vegetative cells. RNA for *nifP* or *nifV* was quantified by RT-qPCR, normalized to *rnpB* (ΔCq), and expressed relative to the control (WT or +N) (ΔΔCq) as shown in the graphs. ***, p <0.001; unlabeled values were not statistically significant.

**Fig. 6.**
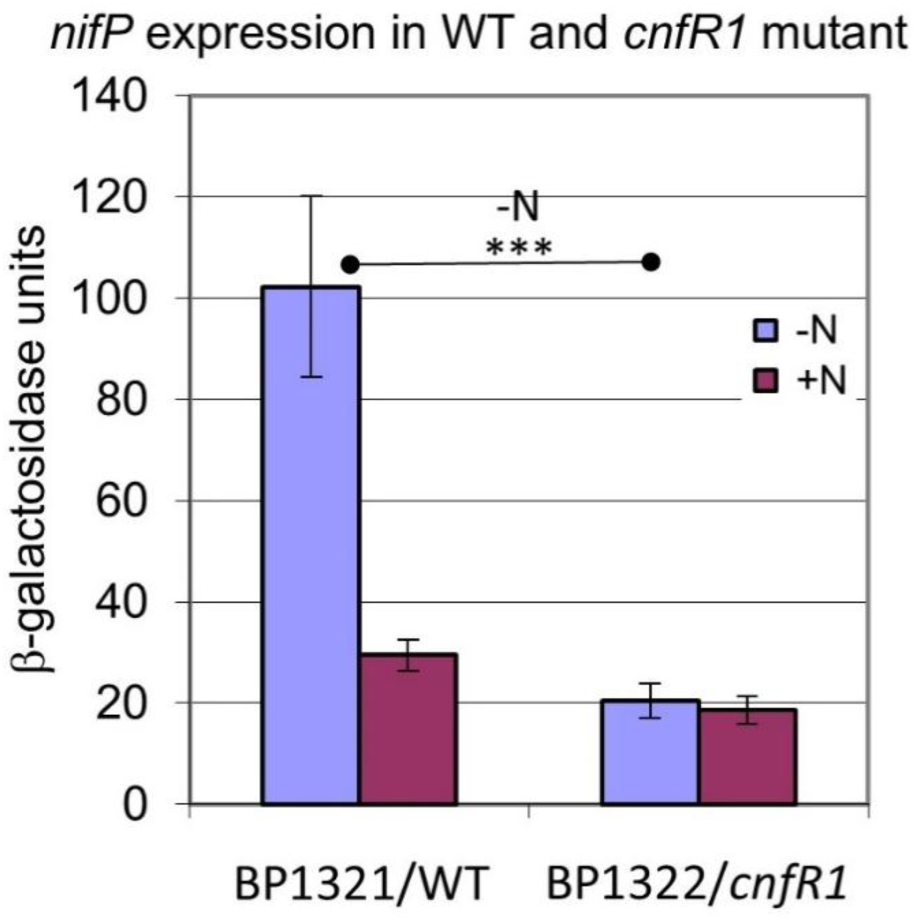
Expression of P*_nifP_*:*lacZ* in WT vs. a *cnfR* mutant. The *nifP* promoter region in the plasmid used to create SM6 (see Fig. 2B) was recombined into the chromosome yielding a full-length *nifP* upstream region fused to *lacZ* at a position 162 bp inside the *nifP* coding region in BP1321 (WT) or BP1322 (a *cnfR1* mutant). β-galactosidase was measured in cells grown +N or after 24 h of nitrogen deprivation (-N). ***, p <0.001; comparisons of unlabeled values were not statistically significant.

Expression of *nifV1* in the WT strain under aerobic conditions was about 34-fold higher in −N growth conditions vs. +N conditions and about 3-fold higher under anoxic −N growth conditions vs. +N conditions (Fig. 5B); however, only the increase under aerobic conditions was statistically significant. In the *cnfR1* mutant, under aerobic −N growth conditions, *nifV1* expression was similar to WT (**Fig. 5B),** indicating that CnfR1 does not control the expression of *nifV1*. In a *cnfR2* mutant, under anoxic −N growth conditions, *nifV1* expression was slightly higher than the WT (**Fig. 5B**), but this was not statistically significant. These data indicate that *nifV1* was expressed only in heterocysts but, unlike most other *nif1* genes, not under the control of CnfR1.

### CnfR1-dependent expression of *nifP* in heterocysts

The apparent increase in *nifP* expression in the WT strain vs. the *cnfR1* mutant (Fig. 5A) suggested that there was greater expression of *nifP* in heterocysts than in vegetative cells and that expression in heterocysts depended on CnfR1. Using a fluorescein-conjugated β-galactosidase substrate, we imaged fluorescent cells expressing *nifP*:*lacZ* (SM5). SM5 cells grown aerobically −N formed heterocysts that fluoresced brightly, indicating that *nifP* was highly expressed in heterocysts (**Fig. 7A–B**). However, consistent with the β-galactosidase assays for SM5 (**Fig. 2B**), vegetative cells in cultures grown with and without fixed nitrogen also fluoresced (**Fig. 7**). Therefore, *nifP* was expressed constitutively at a low level in vegetative cells grown −N or +N, but its expression was greatly increased in heterocysts by CnfR1 activation of the *nifP* promoter.

**Fig. 7.**
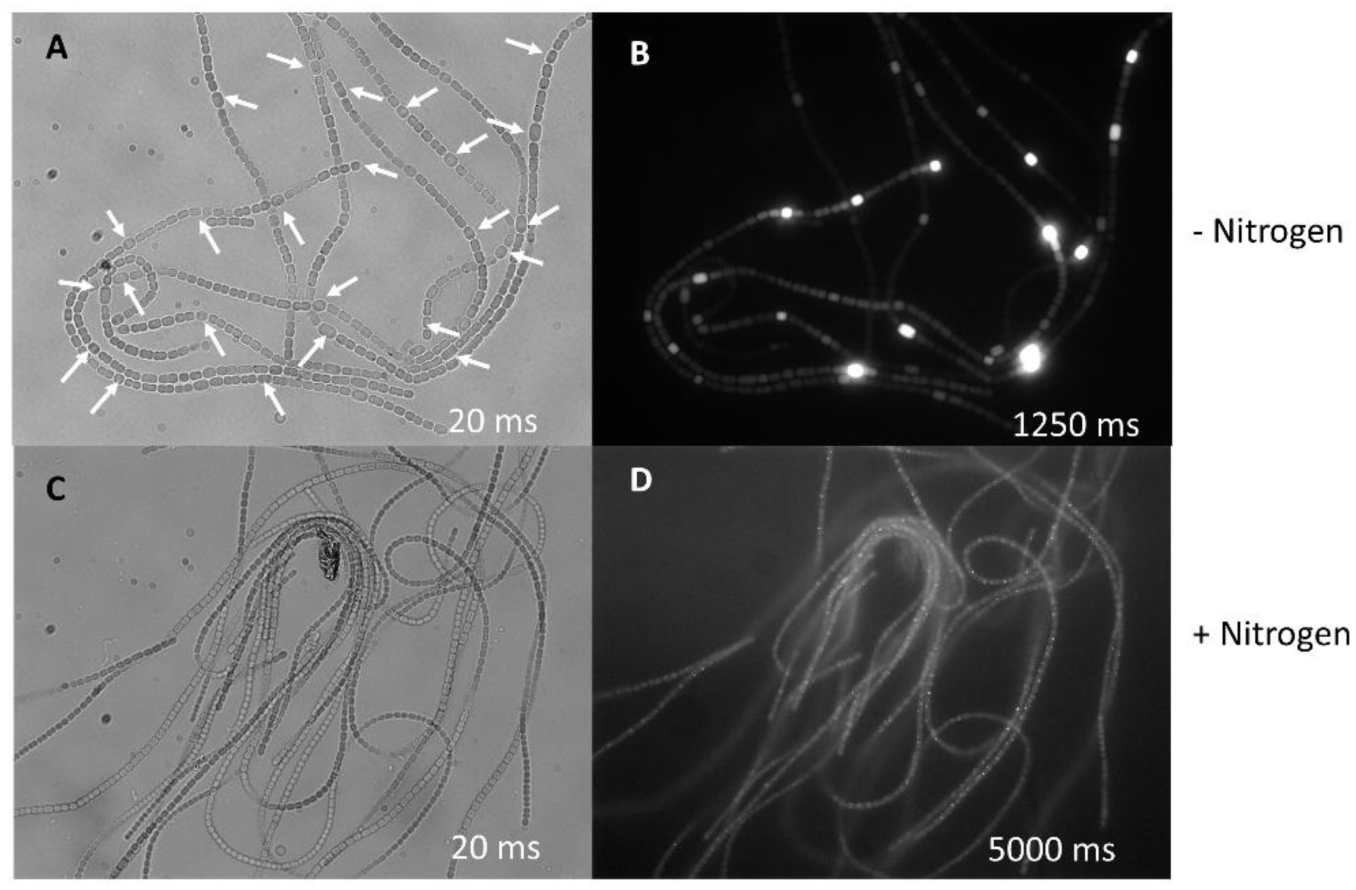
Expression of *nifP* in filaments of *A. variabilis*. SM5 (*nifP*:*lacZ* fusion) was grown without (**A** and **B**) or with (**C** and **D**) fixed nitrogen aerobically for 24 h for heterocyst development. Filaments were visualized by light microscopy (**A** and **C**). β-galactosidase was visualized using a fluorescein □-galactosidase substrate by fluorescence microscopy (**B** and **D**) for the exposure times indicated on each panel. White arrows indicate representative heterocysts. Cells grown +N (**A** and **C**) have no heterocysts.

## Discussion

NifP is serine O-acetyltransferase that converts serine to O-acetyl serine, an intermediate in the synthesis of cysteine, an amino acid used by NifS to create the iron-sulfur clusters in nitrogenase (Evans et al., 1991). The cysteines in the nitrogenase structural proteins combined with those required for the P-clusters, the FeMo-co, and the Fe-S clusters create a high demand for cysteine for nitrogen fixation. While NifP is not required for nitrogen fixation in the heterotrophic proteobacterium *A. vinelandii* (Jacobson et al., 1989a), in the closely related strain *A. chroococcum*, loss of *nifP* slows growth under nitrogen-fixing conditions (Evans et al., 1991). However, in the nonheterocystous cyanobacterium *L. boryana*, loss of NifP reduces nitrogenase activity by 80% compared to WT levels and inhibits diazotrophic growth (Tsujimoto et al., 2014). The *nifP* mutant in *A. variabilis* behaved similarly to the *nifP* mutant in *A. vinelandii* (Jacobson et al., 1989b) with no defect in growth or nitrogen fixation. The genome of *A. variabilis* is annotated to have two additional copies of CysE, suggesting that NifP is redundant and the other two enzymes likely provide sufficient cysteine for nitrogenase synthesis.

Although CnfR1 activates *nifB1* and *nifP*, CnfR2 did not regulate *nifP* as *nifP* was expressed in anoxic vegetative cells in a *cnfR2* mutant. However, *nifP* was also expressed +N; hence, there may be sufficient O-acetyltransferase in anoxic vegetative cells –N to support *nif2*-mediated nitrogen fixation, or *nifP* may be further activated by other transcription factors induced by nitrogen deprivation.

NifV, which produces homocitrate for nitrogenase, is essential for nitrogen fixation in *Klebsiella pneumoniae* and *A. vinelandii* unless cells are provided with homocitrate, which restores the Nif^+^ phenotype (Hoover et al., 1988; Madden et al., 1991). In *Nostoc* PCC 7120, loss of either *nifV1* or *nifV2* decreases nitrogenase and growth, whereas the double mutant virtually abolishes growth and nitrogenase activity. While *nifV1* is expressed only in heterocysts, *nifV2* is expressed constitutively (Masukawa et al., 2007). *A. variabilis* has three copies of *nifV*; two are homologs of *nifV1* and *nifV2* in *Nostoc* sp. PCC 7120; however, the third copy, *nifV3*, is directly upstream of the *vupABC* genes that encode a vanadate transporter required for the synthesis of the V-nitrogenase (Pratte and Thiel, 2006), which is not present in *Nostoc* sp. PCC 7120. Here, we demonstrated that in *A. variabilis*, loss of *nifVZT1*, which is expressed only in the absence of fixed nitrogen, had no effect on nitrogen fixation, likely because the second and/or third copy of the *nifV* gene in this strain provides sufficient homocitrate for nitrogenase activity. The *nifVZT1* operon is just downstream of *nifP*, but has its own promoter and, unlike *nifP*, these genes were not expressed in anoxic vegetative cells grown –N. Unlike the major *nif1* and *nif2* clusters, expression of *nifVZT1* was not activated by CnfR1, consistent with the absence of the conserved motifs found upstream of *nifB1*, *nifB2*, and *nifP*.

The conserved motifs upstream of the *nifB1*, *nifB2*, and *nifP* promoter regions in *A. variabilis* (Vernon et al., 2017) and the *nifB-nifP* intergenic region in *L. boryana* (Tsujimoto et al., 2016) (see Fig. 1A) have different roles depending on the gene. Here, we demonstrated that motifs 1B and 2B were important for the regulated transcription of *nifP*, with 2A having a lesser role. Upstream motif 1B inhibited transcription of *nifP* regardless of nitrogen availability; thus, it is likely that this inhibition occurs in vegetative cells. In contrast, in *nifB1*, we showed that loss of either 1A or 1B results in about a 50% decrease in transcription of *nifB1* with a further decrease when both motifs are mutated (Vernon et al., 2017). Further, although the conserved regions upstream of *nifB1* and *nifB2* in *A. variabilis* are similar, the replacement of the *nifB2* conserved region with the similar region from *nifB1* results in complete loss of *nifB2* expression (Vernon et al., 2017). Differences in the effects on transcription for these conserved *nifB*/*nifP* motifs are also evident for the filamentous, non-heterocystous strain, *L. boryana.* The promoter region of the *L. boryana nifB* gene was fused to *luxAB* in the heterologous cyanobacterial host, *Synechocystis* sp. PCC 6803, and *cnfR* was expressed there under the control of the *trc* promoter. In this system, modification of the sequences corresponding to 2A or 2B (named IV-V and VI-VII in *L. boryana*) abolishes transcription by CnfR, whereas modification of 1B (named I-II in *L. boryana*) has little effect on transcription (Tsujimoto et al., 2016). DNA fragments from *Nostoc* sp. PCC 7120 containing these conserved regions upstream of *nifB*, *nifP*, and *alr2522*, a gene of unknown function, bind *purified* recombinant CnfR in vitro (Rachedi et al., 2023).

In *A. variabilis*, in addition to the conserved *nifB2* 5′ untranslated region, expression of *nifB2* requires sequences within the 5′ end of the *nifB2* gene (Vernon et al., 2017). Similarly, here we found that *lacZ* fusions that extended into the coding region of the *nifP* gene provided better nitrogen-controlled transcription than fusions lacking these extensions. However, as *nifP* was expressed constitutively in vegetative cells, the coordination of expression in heterocysts vs. vegetative cells likely involves other factors such as NtcA. For example, normally, the expression of *nifB2* requires NtcA because the expression of its activator, CnfR2, is NtcA dependent (Pratte and Thiel, 2016). Although *nifB2* can be activated by CnfR1 when it is expressed in vegetative cells from a heterologous Co-inducible *coaT* promoter, that expression requires NtcA, suggesting that NtcA may have a role in the expression of *nif* genes that are expressed in vegetative cells (Pratte and Thiel, 2021). However, the varied effects on gene expression of modifications to the promoter regions of different *nif* genes, particularly those examined by promoter fusions to reporter genes, make the elucidation of the role of these regions in the regulation of gene expression difficult to interpret.

## Materials and Methods

### Strains and growth conditions

Details of growth conditions for *A. variabilis* FD have been described previously (Pratte and Thiel, 2016). Briefly, cells were in an 8-fold dilution of Allen and Arnon (AA) medium supplemented, when appropriate, with 5 mM NH_4_Cl, 10 mM *N*-tris (hydroxymethyl)methyl-2-aminoethanesulfonic acid (TES), pH 7.2 at 30°C, with light at 100 to 120 μE m^−2^ s^−1^. For determination of specific growth rate during true exponential growth, freshly grown cultures in AA/8 were diluted to an OD_720_ of <0.05 and grown in quadruplicate to a final maximum OD_720_ of about 0.15 in shallow 3 ml-well plates to minimize self-shading of the cells. The OD_720_ of culture samples was measured every 4– 6 h, and the growth rate was determined during early exponential growth before light limitation yielded linear growth.

### Construction of strains

Plasmids constructed here are described in Table 1 below, and primers are shown in Table 2 below. Plasmid inserts were verified by sequencing. All plasmids were conjugated to the WT or mutant strains indicated in the strain descriptions (Table 1) as described previously (Elhai and Wolk, 1988) and selected with the appropriate antibiotics.

**Table 1.**
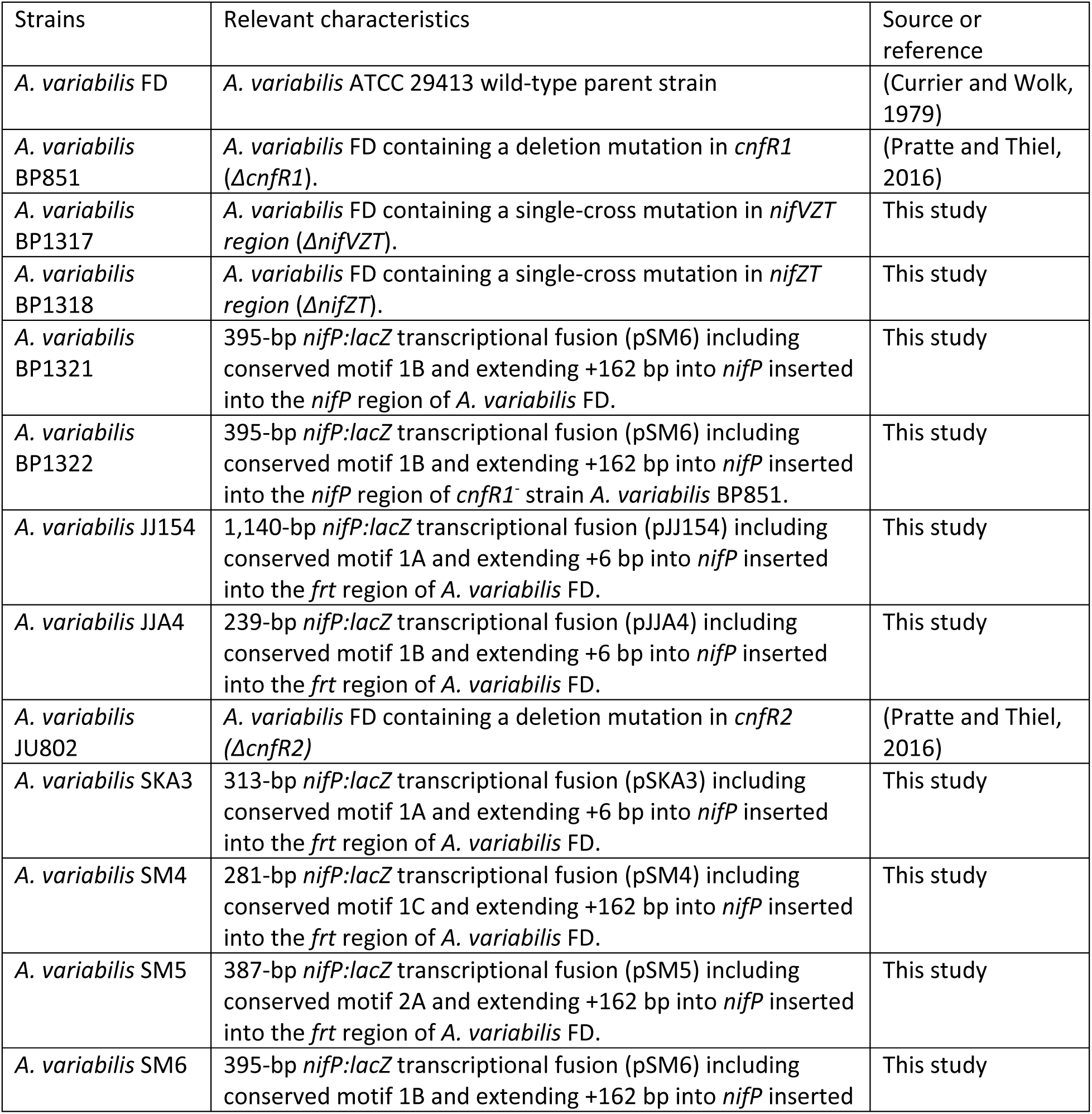
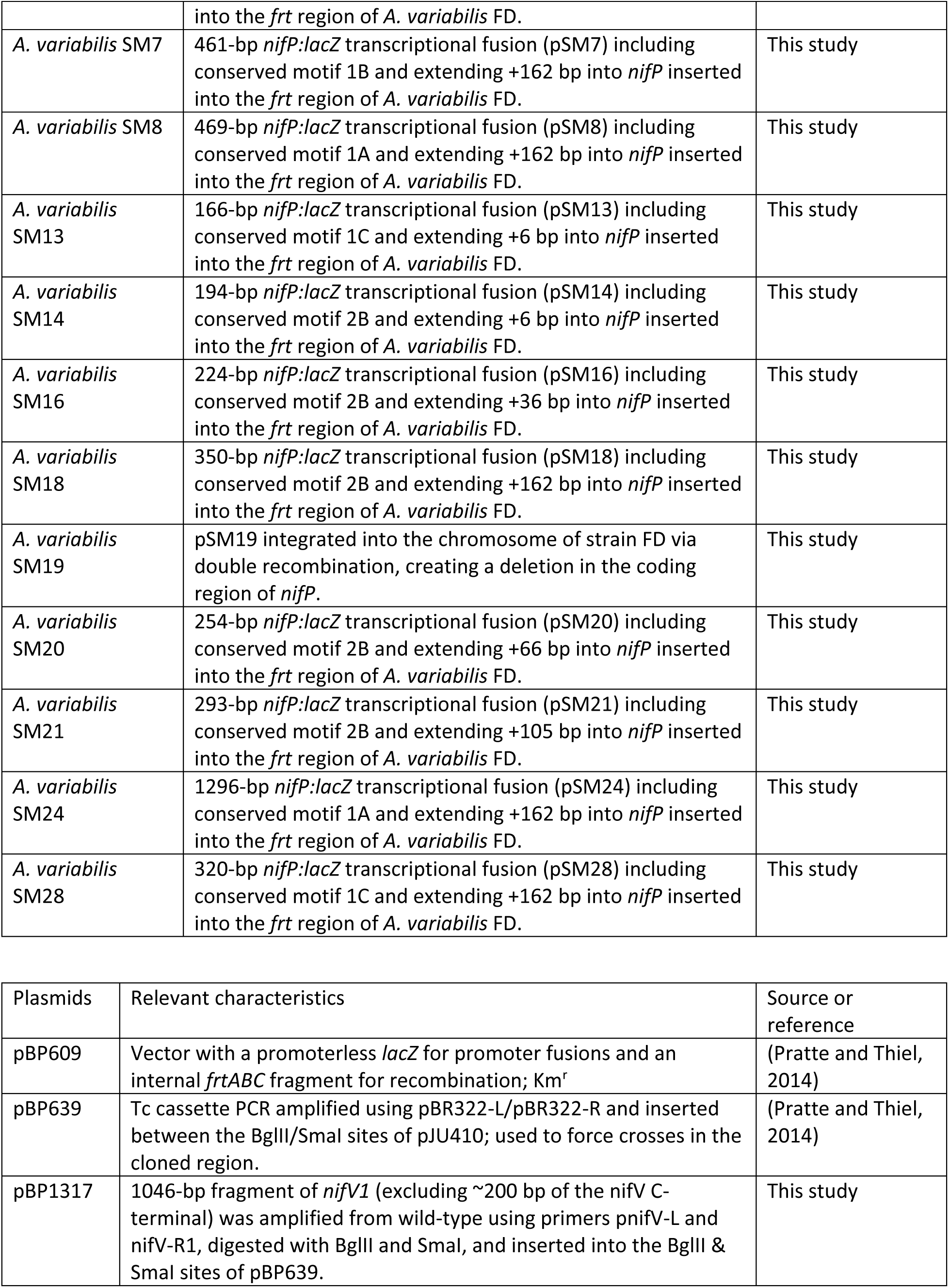
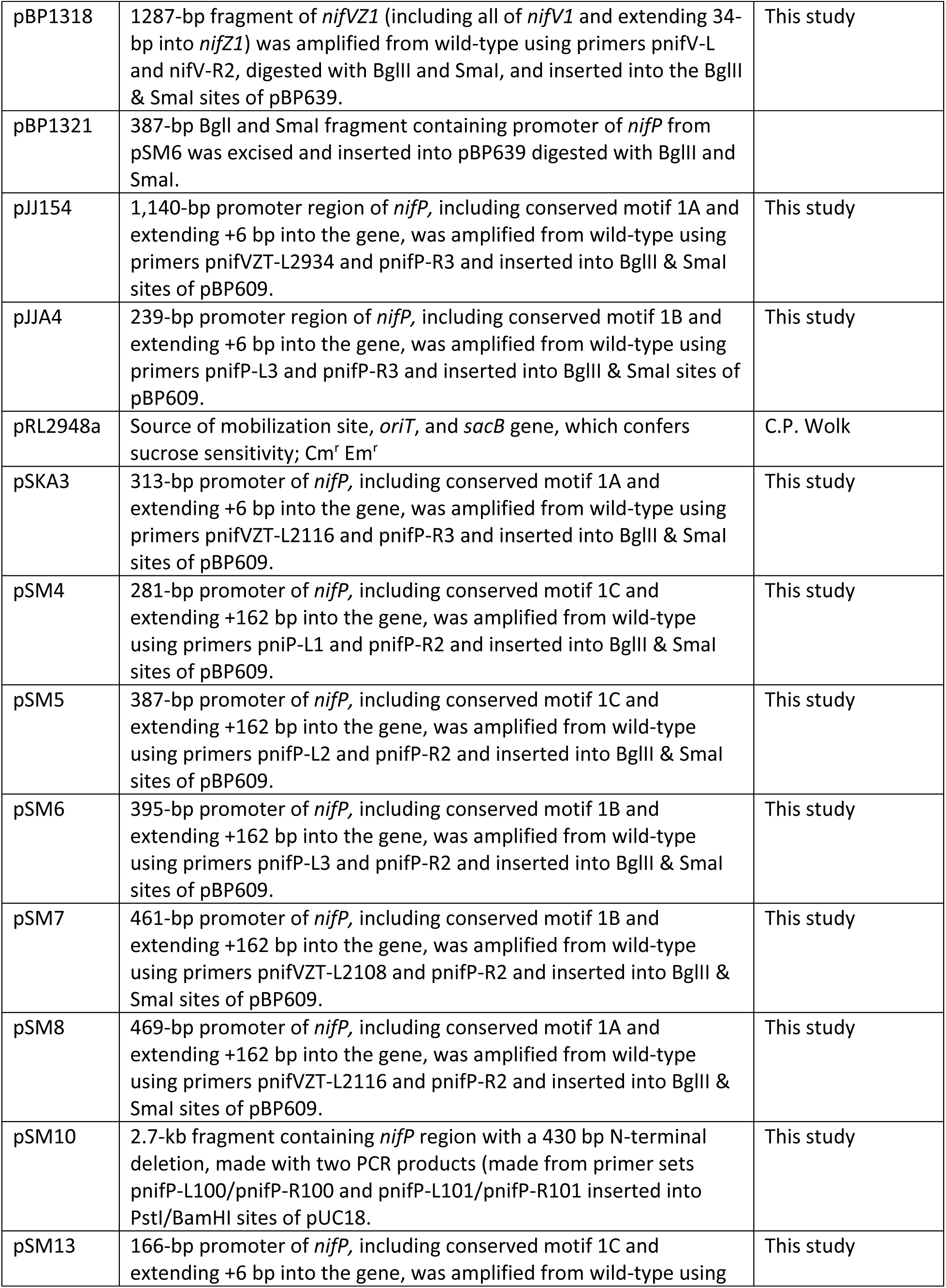
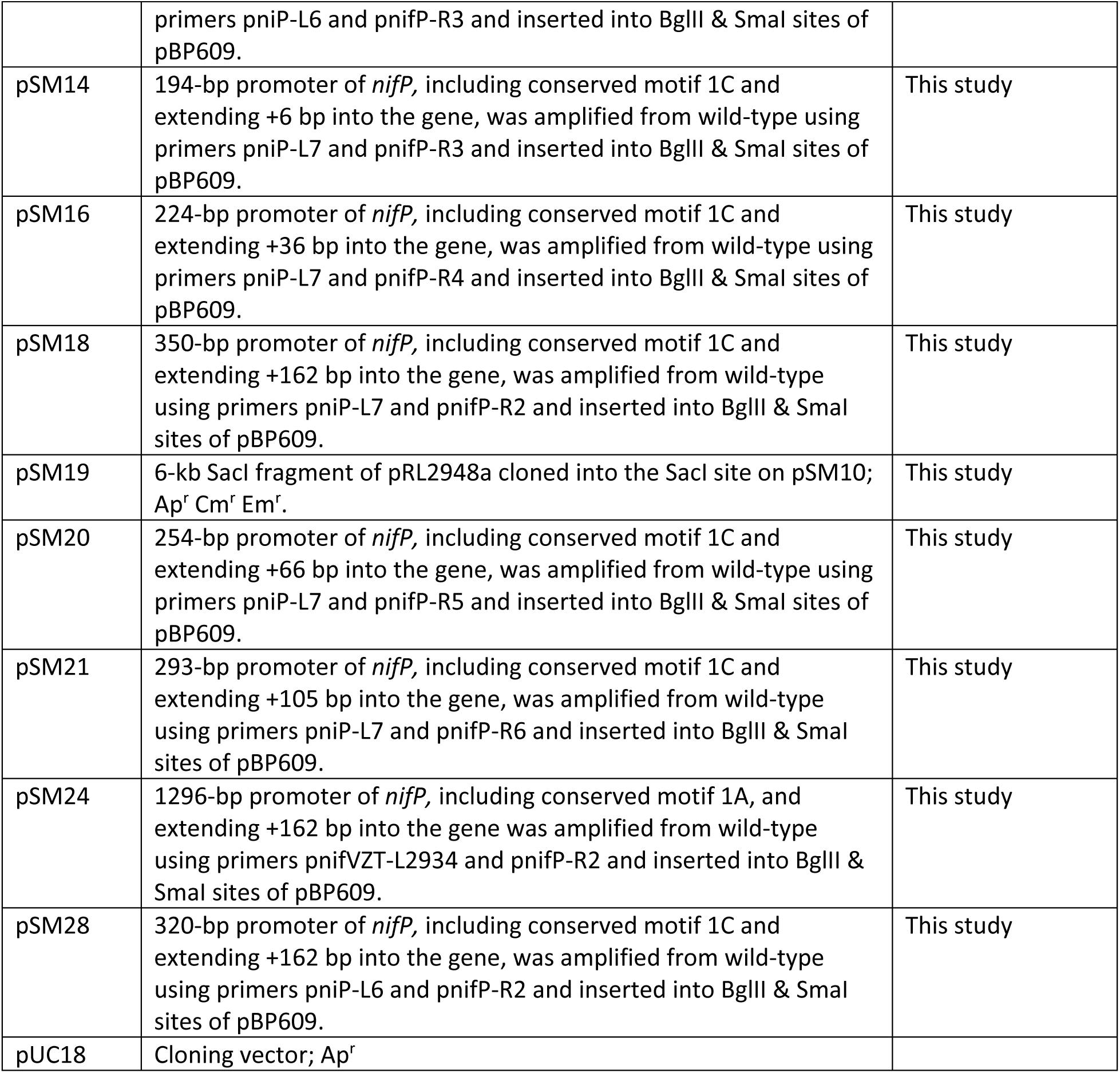
Strains and plasmids.

**Table 2.**
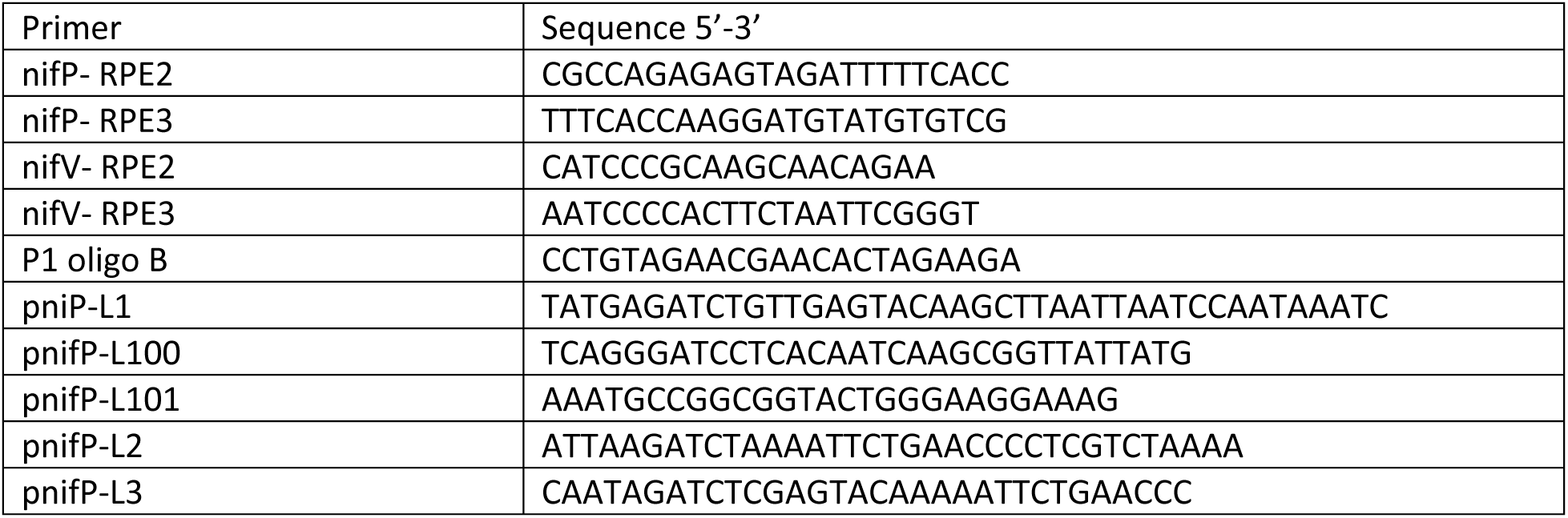
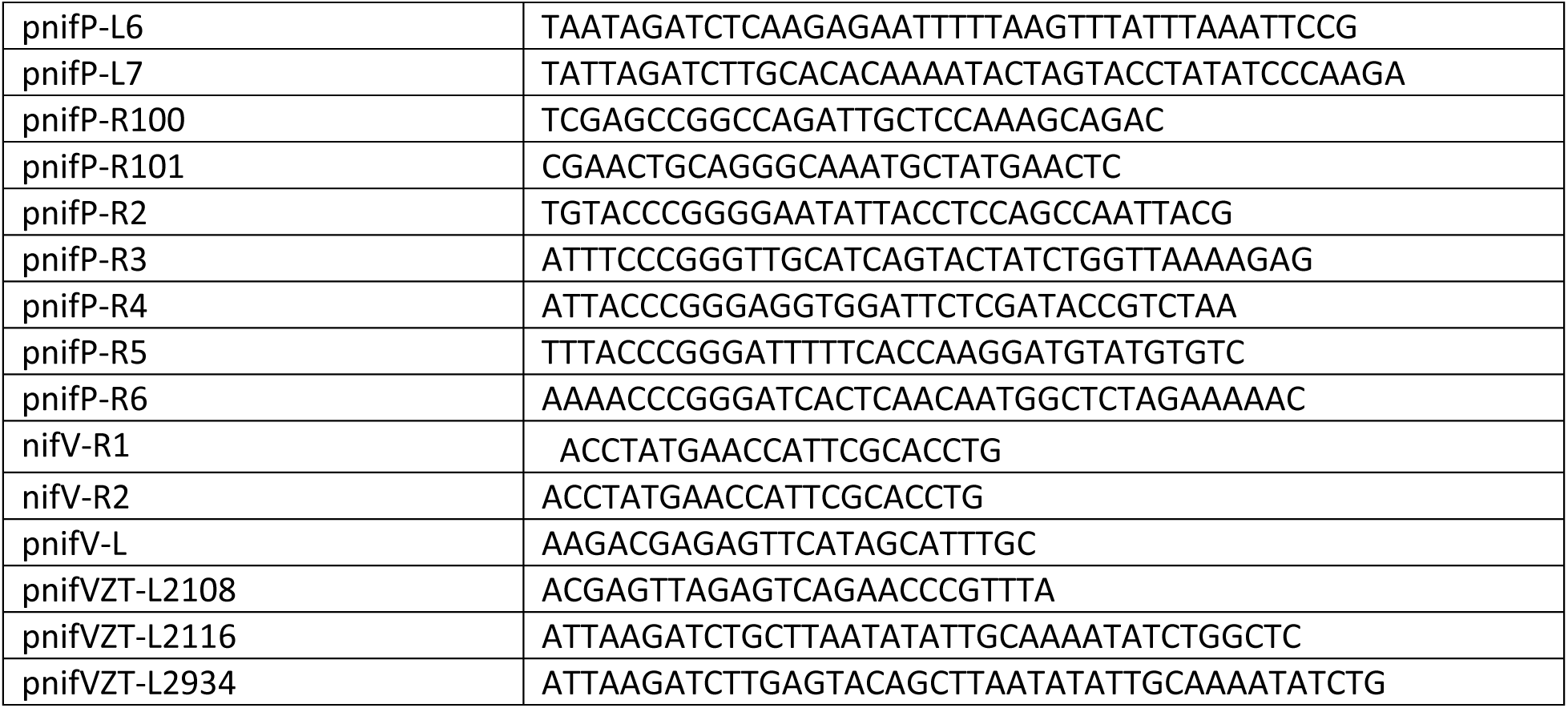
Primers.

### nifP:lacZ fusions

Promoter fragments of various lengths upstream of *nifP* and/or into the 5’ end of the *nifP* gene were PCR amplified using the primers listed for each plasmid in Table 1 and inserted between the BglII/SmaI sites in the vector pBP609 to create various P*_nifP_*:*lacZ* fusions that were conjugated into WT strain FD. Single recombinants that had P*_nifP_*:*lacZ* fusions were first selected as Nm^R^Sm^R^Sp^R^ and then screened for exconjugants that recombined in the *frtBC* region, which were unable to grow heterotrophically on fructose in the dark (Ungerer et al., 2010). In contrast, strains BP1321 and BP1322 have a full-length length *nifP* upstream region fused to *lacZ* fusion expressed in a WT (BP1321) or a *cnfR* mutant (BP1322) background. Plasmid pSM6 (Table 1) with a 395-bp *nifP:lacZ* transcriptional fusion including conserved motif 1B and extending +162 bp into *nifP* was recombined into the *niP* region of WT (BP1321) or *cnfR1* mutant (BP1322) chromosome yielding a full-length *nifP* upstream region fused to *lacZ* at a position 162 bp inside the *nifP* coding region.

### nifP, nifV1 and nifZT1 mutants

The *nifP* mutant SM19 was created by double recombination of plasmid pSM19 into the chromosome of strain FD, creating a 430 bp N-terminal deletion in the coding region of *nifP*. The *nifP* deletion was created in plasmid pSM10 (Table 1) by fusing two PCR products (made from primer sets pnifP-L100/pnifP-R100 and pnifP-L101/pnifP-R101 into PstI/BamHI sites of pUC18. A 6-kb SacI fragment of pRL2948a was cloned into the SacI site on pSM10, making the mobilizable plasmid pSM19 that was conjugated into *A. variabilis* and *sacB* screening for sucrose sensitivity was used to obtain a double recombinant. These strains and plasmids are described in Table 1 below.

BP1321 and BP1322 have a P*_nifP_*:*lacZ* fusion expressed in a WT (BP1321) or a *cnfR* mutant (BP1322) background. Plasmid pSM6 (Table 1) with a 395-bp *nifP:lacZ* transcriptional fusion including conserved motif 1B and extending +162 bp into *nifP* was recombined into the WT (BP1321) or *cnfR1* mutant (BP1322)chromosome yielding a full-length *nifP* upstream region fused to *lacZ* at a position 162 bp inside the *nifP* coding region.

A *nifVZT* mutant lacking ∼200 bp of the C-terminus of *nifV* and all of *nifZT1* was made by selection for single-crossover recombination of pBP1317 (described in Table 1) into the WT chromosome. Because the *nifV* fragment in pBP1317 lacks the transcription start site and promoter region of *nifVZT1*, BP1317 recombinants have one *nifVZT1* copy that lacks the *nifVZT1* promoter but has the structural genes and one copy that has the promoter region but lacks the C-terminus of *nifV1* and lacks all of *nifZT1*. Similarly, the *nifZT1* mutant BP1318 was constructed by single-crossover recombination of pBP1318 (described in Table 1) into the WT chromosome. The pBP1318 *nifV* region 1) lacks the transcription start site and promoter region of *nifVZT1*, 2) has a complete *nifV1* gene, and 3) lacks *niZT1* genes. Therefore, BP1318 recombinants have one copy of *nifVZT1* that lacks the *nifVZT1* promoter but has the structural genes and one copy that has the promoter region and *nifV1* but lacks *nifZT1*.

### Nitrogen step-down experiments

Cultures were freshly grown in AA/8 with 5 mM NH_4_Cl and 10 mM TES, pH 7.2, diluted 1:100 in the same medium, and grown to OD_720_ = 0.1–0.2. Cells were washed three times in AA/8, diluted to OD_720_ = 0.1 at 0 time, and grown with light and shaking at 30°C as described above, with or without 5 mM NH_4_Cl and 10 mM TES, for 24 h for assays.

### RNA Isolation, RT-qPCR, and 5′ RACE (rapid amplification of cDNA ends)

RNA isolation and RT-qPCR have been described (Pratte and Thiel, 2016). Briefly, 40 ng of purified DNA-free RNA was converted to cDNA using iScript Reverse transcription Supermix for RT-qPCR (BioRad), and 0.8 ng cDNA in a 10 μl reaction mixture was amplified by PCR using 5 pmol of gene-specific primers and 1x SsoAdvanced SYBR Green supermix (BioRad). The Cq values were normalized to the housekeeping gene, *rnpB*.

5′ RACE was performed as we described previously (Pratte and Thiel, 2016). Briefly, purified DNA-free RNA was treated with RNA 5′ polyphosphatase (Epicentre, Madison, WI), and an RNA adapter, RNAoligo09 (Pratte and Thiel, 2016), was ligated using T4 single-stranded RNA ligase (NEB). The ligated RNA was reverse transcribed using Superscript III (Invitrogen) using the following primer for *nifP,* nifP-RPE2, and for *nifV*, nifV-RPE2. PCR was performed using the left primer P1 oligo B and for *nifP*, nifP-REP3, and for *nifV*, nifV-RPE4. The PCR band was excised and sequenced to determine the 5′ end. Comparison with RNA samples not treated with RNA 5′ polyphosphatase served to distinguish primary vs. secondary processed transcripts.

### Acetylene reduction assays

Assays are described in (Pratte and Thiel, 2016). Briefly, acetylene was injected into 5-ml cultures in 16-ml sealed Hungate tubes that were incubated in the light as described above for 60 min. We measured ethylene using a Shimadzu gas chromatograph with a 6-foot Poropak N column at 75°C.

### β-Galactosidase assays

The assays are described in detail in (Vernon et al., 2017). Briefly, using cultures grown as described above for nitrogen step-down experiments, three 2-ml biological replicates of each culture were grown for 24 h at 30°C with shaking and illumination at 100 to 120 µE m^−2^ s^−1^ in 12-well microtiter plates. In order to normalize b-galactosidase activity, an OD720 measurement of all cultures was determined prior to performing β-galactosidase assays. β-galactosidase assays were performed for 10 min at 30_°_C using 0.5 M NaPO4, pH 7.4, 1.0% sarkosyl, and 1.6 mg/ml ortho-nitrophenyl-β-galactoside (ONPG) in 96-well flat bio-assay microtiter plates with 250 µl of sample. We used three biological replicates, with quadruple technical replicates for each biological replicate. After the reaction was stopped, OD_420_ and OD_665_ measurements were used to 1) determine ONP (OD_420_), 2) to correct for chlorophyll and light scattering from permeabilized cells (OD_665_), and 3) to calculate β-galactosidase activity as 1000 X ((OD_420_ – (1.58 X OD_665_))/(OD_720_ X time of assay in min)) (Vernon et al., 2017).

### In situ localization of β-galactosidase

The procedure is described in detail in (Vernon et al., 2017). Briefly, cells that were grown as described for β-galactosidase assays were fixed for 15 min at 25°C with 0.04% glutaraldehyde, washed free of glutaraldehyde, and incubated in the dark at 37°C for 30 min with 100 µM 5-dodecanoyl-aminofluorescein di-β-D-galactopyranoside in 25% dimethyl sulfoxide. Washed cells were resuspended in an anti-fade mounting medium, and expression of *lacZ* in cells was visualized on a Zeiss Confocal microscope using fluorescein excitation and emission wavelengths to detect β-galactosidase and rhodamine excitation and emission wavelengths to visualize native cyanobacterial autofluorescence from phycobiliproteins.

### Statistical analysis

Data are shown as the mean ± standard deviation. The significance of the differences between the means for two values was analyzed using an unpaired, two-tailed Student’s *t*-test, with *P* values of < 0.05 considered statistically significant.

## Data availability

All processed data (e.g., relative gene expression and β-galactosidase activity) used in the experiments are provided here. Additional raw data may be requested from the authors.

## Acknowledgment

This work was supported by the National Science Foundation grant MCB-1818298.

